# Large-Scale Evolutionary Patterns of Protein Domain Distributions in Eukaryotes

**DOI:** 10.1101/142182

**Authors:** Arli A. Parikesit, Peter F. Stadler, Sonja J. Prohaska

## Abstract

The genomic inventory of protein domains is an important indicator of an organism’s regulatory and metabolic capabilities. Existing gene annotations, however, can be plagued by substantial ascertainment biases that make it difficult to obtain and compare quantitative domain data. We find that quantitative trends across the Eukarya can be investigated based on a combination of gene prediction and standard domain annotation pipelines. Species-specific training is required, however, to account for the genomic peculiarities in many lineages. In contrast to earlier studies we find wide-spread statistically significant avoidance of protein domains associated with distinct functional high-level gene-ontology terms.

**1998 ACM Subject Classification** J.3 Life and Medical Sciences

## 1 Introduction

Proteins embody a wide variety of functions in a cell, ranging from enzymatic activity to structural scaffolding. The range of an organism’s biochemical capabilities, both metabolic and regulatory, is thus largely encoded in its protein content. This is true even though RNA-based mechanisms can play a fundamental role as in the case of post-transcriptional regulation by microRNAs. In fact, the presence or absence of RNAi pathways, for instance, can be inferred from the presence or absence of its protein components [8]. Large-scale trends in evolution such as an increased complexity of transcriptional regulation [16, 28], the diversification of chromatin modification [25], or novel modes of post-transcriptional processing are visible in comparisons of the predicted protein complements and thus are focal features of most genome papers.

Most proteins are composed of smaller building blocks. A protein domain typically forms a compact three-dimensional structure that is frequently stable and foldable on its own and conveys a specific molecular function such a particular catalytic activity or binding specificity. Protein domains are characterized by local amino-acid patterns and hence can be annotated computationally in protein sequences. Several databases, most notably `Pfam` [26] and `SUPERFAMILY` [7], provide large collections of domain descriptions in the form of Hidden Markov Models HMMS for this purpose. Since protein domains are also regarded as functional units, the same databases provide maps to link domains with `GeneOntology` (GO) terms. As GO terms are primarily associated with entire proteins, these maps are obtained at least in part computationally [7, 27]. Conversely protein function can be computed from domain content [10].

Protein domains also constitute units in evolutionary terms. They can be readily recombined in different arrangements leading to proteins that utilize different combinations of the same (types of) molecular interactions to fulfill different higher-level functions [1, 22, 4]. Over very large evolutionarily time scales, such as those of interest in a comparative analysis of the eukaryotic kingdoms, it thus becomes impossible in many cases to identify orthologous proteins since larger proteins more often than not a composite of domains deriving from several ancestral sources [18, 14]. Fusions, fissions, and terminal loss have turned out to be much more frequent than the innovation of novel protein domains [3, 37]. The abundance and co-occurrence of domains thus becomes the most natural and promising framework to understand patterns of protein evolution at kingdom-level time-scales, see e.g. [13, 25, 35]. In [37], for instance, showed that frequent gains and losses of domains lead to significant differences in functional profiles of major eukaryotic clades. Their results argue for a complex last eukaryotic common ancestor and reveal suggest that animals are gaining increased regulatory complexity at the expense of their metabolic capabilities. Similarly, the rise of chromatin-based regulation mechanisms in crown-group eukaryotes can be traced by considering abundances and co-occurrences of the relevant protein domains [25].

The most complete information on the protein complement can be inferred from the genome sequence. In fact, only two thirds of the predicted human proteins have been directly observed so far [21]. For most of the less-studied species, on the other hand, the set of predicted proteins in the current genome annotations is far from complete. For example, the number of annotated transcripts varies by more than a factor of three even between great ape genomes [24]. The accumulation of transcriptomics data in a few well-studied organisms such as human, mouse, or fruit fly, on the other hand, leads to an increasing number of annotated splice variants and transcripts with alternative start sites, and thus to an increasing number of redundant protein variants. In our previous studies we have argued, therefore, that the large ascertainment biases in present day protein databases make it effectively impossible to use these data for a quantitative comparison of protein domain abundances across species [24, 23]. Instead, we proposed to use *de novo* gene predictions to obtain quantitatively comparable estimates, Fig. 1, and showed that a simple general-purpose gene finder such as genscan [5, 6] already yields plausible numbers.

**Figure 1.**
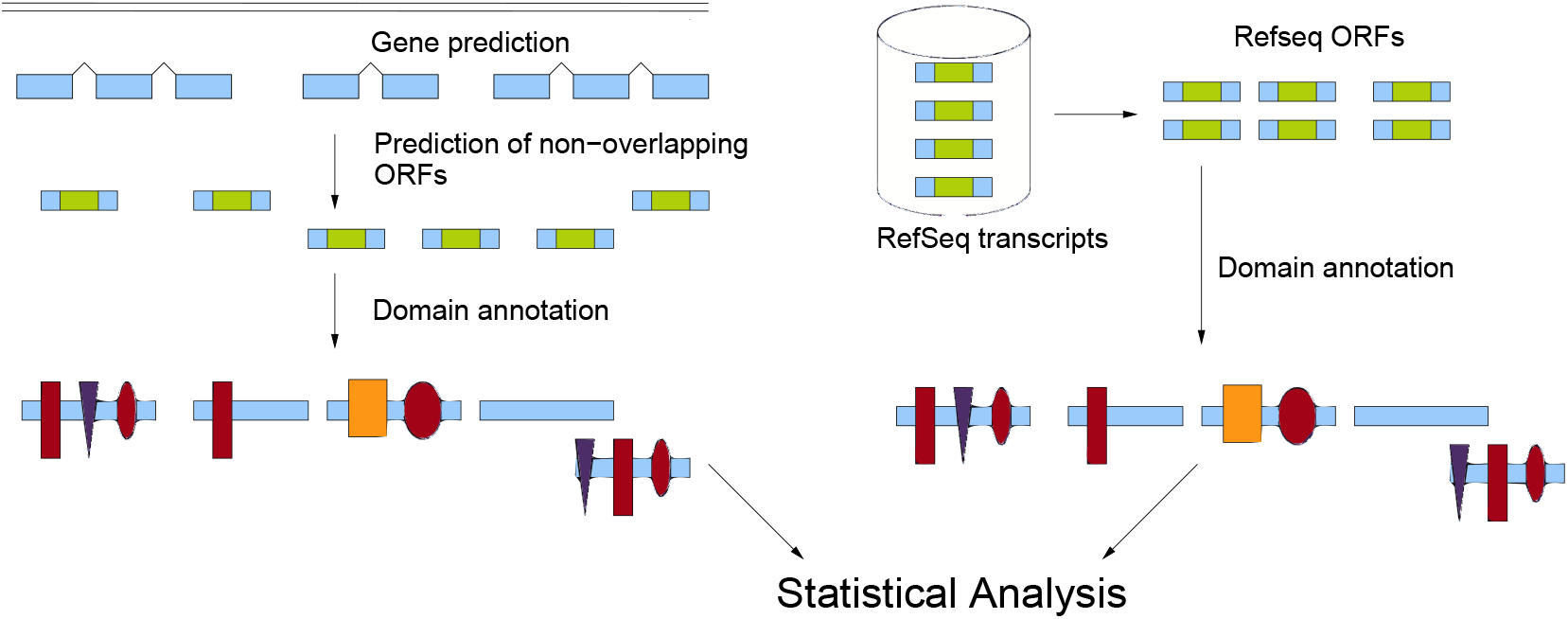
Work flow for the estimation of domain abundance data. We start with a *de novo* gene annotation (l.h.s), here using `AUGUSTUS` to obtain a collection of non-overlapping protein predictions that is as unbiased as possible. Most studies instead start from protein databases that suffer from a variety of ascertainment biases. Protein domains from the `Pfam` or `SUPERFAMILY` database are mapped to the known or predicted proteins and form the basis for subsequent statistical analysis.

Several major lineages of the Eukarya feature gene structures and a genomic organization that is very different from the situation in animals, fungi, or plants. Both *Giardia lamblia* and *Trichomonas vaginalis* are extremely intron-poor; *Trichomonas vaginalis* in addition features very large numbers of paralogs. Kinetoplastids ( *Trypanosoma* and *Leishmania*) produce large polycistronic transcripts from which individual mature mRNAs are produced by trans-splicing, cis-splicing, and polyadenylation [17, 34]. Trans-splicing is also prevalent in the nematodes, but absent from most other animal genomes. Intron-sizes differ dramatically between invertebrates and vertebrates, where intron-sizes of more than 10 kb are not at all uncommon. Another problem is posed by the extreme sequence composition as in the AT-rich genome of *Plasmodium falciparum* [15].

In this contribution we therefore employ `AUGUSTUS` [31] a gene prediction tool that can be adapted to the individual genomes and their peculiarities. In extension of our earlier work we furthermore use both `SUPERFAMILY` and Pfam database of domain annotation.

## 2 Material and Methods

### 2.1 Genome Sequences and Gene Prediction

We consider the following 18 species with sequenced genomes covering the entire phylogenetic range of the eukaryotes: *Homo sapiens* hg19, *Drosophila melanogaster* BDGP5.13, *Caenorhabditis elegans* WS200, *Schizosaccharomyces pombe* EF1, *Aspergillus niger* CADRE, *Arabidopsis thaliana* TAIR9.55, *Clamydomonas* Chlre4, *Tetrahymena* tta1 oct2008, *Plasmodium falciparum* PlasmoDB-7.0, *Leishmania major* Lmj_20070731_V5.2, *Giardia lamblia* WBC6, *Trichomonas vaginalis* TrichDB-1.2, *Trypanosoma brucei* Tb927_May08_v4, *Naegleria gruberi* Naegr1, *Thalassiosira pseudonana* Thaps3, *Phytophthora ramorum* Phyra1_1, *Oryza sativa* OSV6.1, *Dictyostelium discoideum* DDB. Sources are listed in the Supplement http://www.bioinf.uni-leipzig.de/supplements/12-007.

We decided to use AUGUSTUS [32, 31, 30] for gene prediction because the package has gained popularity in genome annotation projects and because it can be trained for applications to a given genome with known cDNAs. For our analysis, we used both “off-line” (local) and “on-line” (web-based) trained models prepared as described in the `AUGUSTUS` tutorial [29]. For the several species, the default training sets are provided at the `AUGUSTUS` website. For these, there is no difference between local and web-based training. For the remaining species, we used the cDNAs available in GenBank. Redundancies were removed with `duplicate.pl` script. The FASTA sequences and their headers were cleaned from meta-characters and gaps. Models were trained both “off-line” and using the pipeline offered at the `AUGUSTUS` website. For our applications, `AUGUSTUS` was configured to generate only non-overlapping protein-coding genes. The predicted protein sequences are part of the `AUGUSTUS` output. We verified with `bed-tools` that no overlapping sequences were contained in the output.

The results of `AUGUSTUS` are compiled in Table 1 together with the `RefSeq` (release 53) genes for each of the 18 species. In order to compare the two training modes of `AUGUSTUS` with each other and with the `RefSeq` annotation we computed their overlaps with `bed-tools` and used `lucidchart` to compute Venn diagrams so that the displayed overlaps in Fig. 2 are drawn to scale.

**Figure 2.**
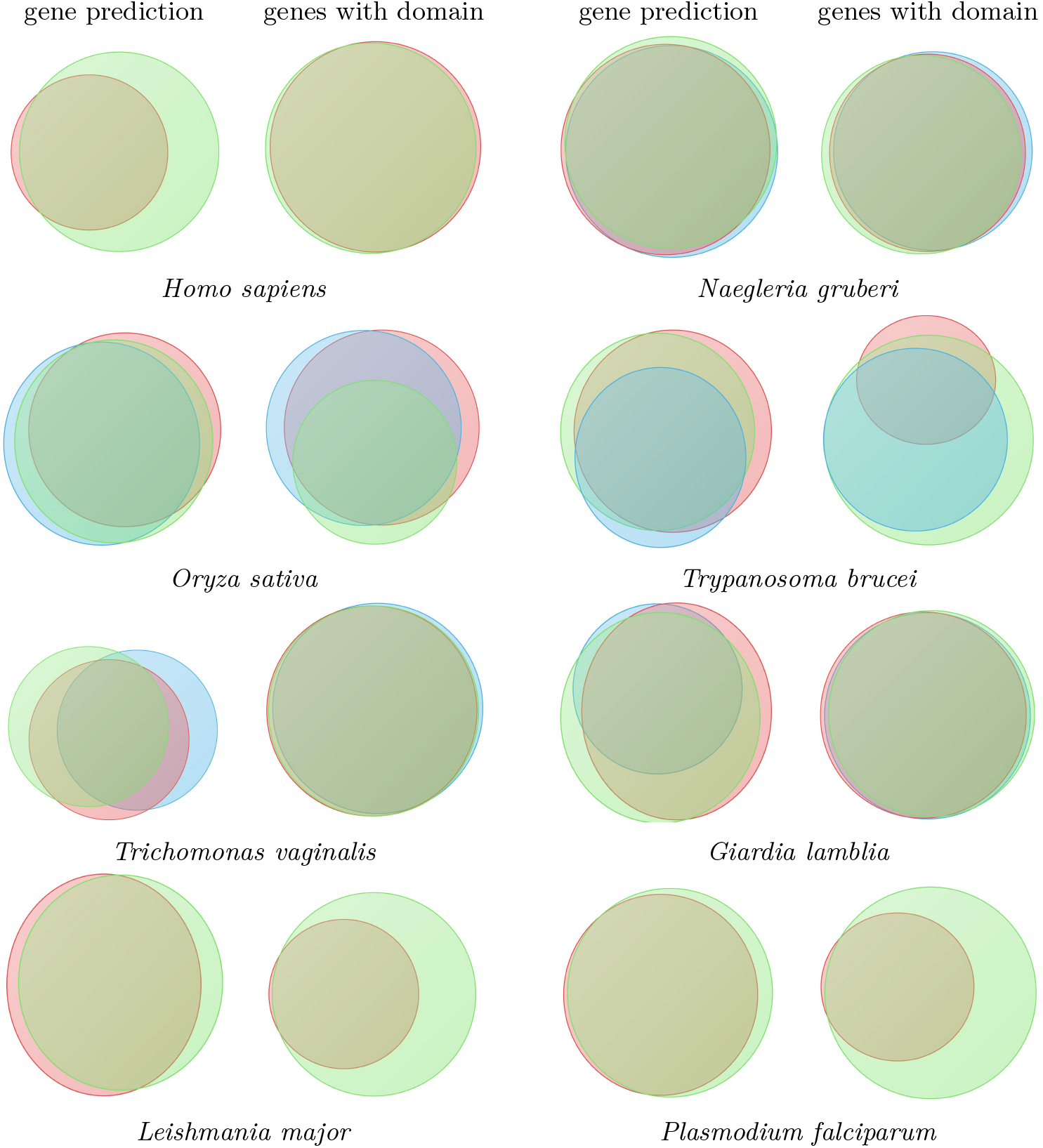
Comparison of gene predictions for 8 of the 18 species. (See online supplement for the remaining data. For each species we show a Venn diagram for both the raw output of the gene predictions and for the subset of proteins with at least one matching `Pfam` model. `RefSeq` is shown in red `AUGUSTUS` prediction with online and offline trained models are shown in blue and green, respectively.

**Table 1.**
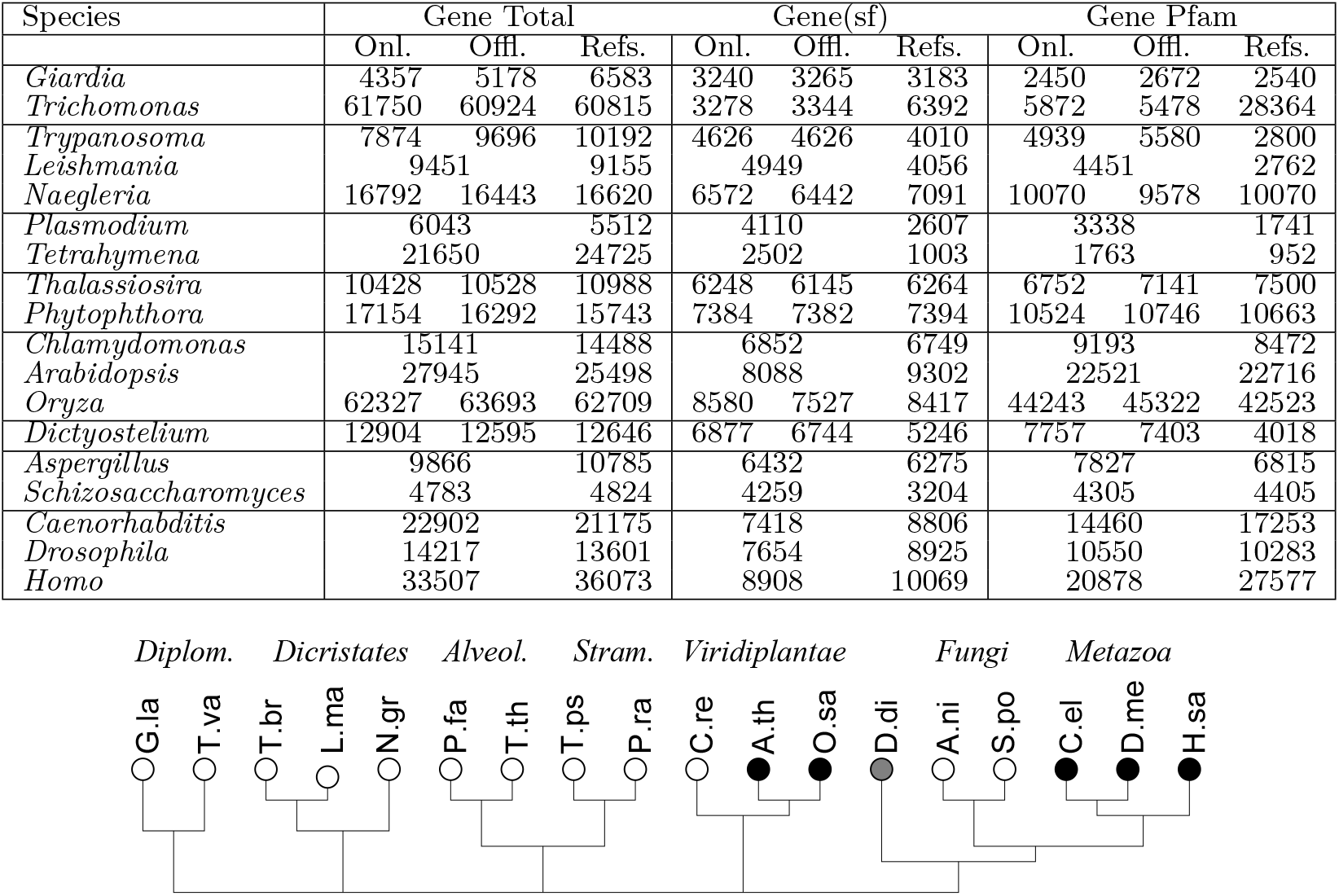
Summary of gene and domain annotation. The first block gives the results from `AUGUSTUS` with both training methods where available and the contents of the `RefSeq` database. The following blocks of columns list the numbers of genes that have at least one `SUPERFAMILY` or `PFAM` domains, respectively. Below, the phylogenetic distribution of the 18 investigates species is summarized [2]. See sect. 2.1 for full species names.

### 2.2 Domain Annotation

We used the entire `Pfam` version 26.0 database, comprising 33672 domain models as well as the entire collection of 9821 Hidden Markov Models (HMMs) provided by the SUPERFAMILY database (version 1.75). In both cases we used `HMMER3.0rc1` [9] with an *E*-value threshold of *E* ≤ 10^−3^ to map the HMMs to the predicted amino acid sequences as well as the `RefSeq` proteins.

In order to test the quality of gene predictions we compared the sub-collections protein sequences with at least one mapped `Pfam` domain between the gene prediction methods and `RefSeq` database. A representative selection of these results is shown in Fig. 2. Overall, the online-trained `AUGUSTUS` predictions have the best coverage of the manually curated `RefSeq` and are hence used as data basis for subsequent quantitative analysis.

### 2.3 Functional Classification

The domain databases contain thousands of distinct domain models. Few domains thus appear a sufficiently large number of time to allow a quantitative statistical analysis of their occurrences. Thus we pool the data by functional categories. The `SUPERFAMILY` database offers a “Structural Domain Functional Ontology” providing functional and phenotypic annotations of protein domains at the ***superfamily*** and ***family*** levels [7]. The `Pfam` annotation is already integrated into GO database, providing a mapping from `Pfam` domains to GO ontology terms [33, 26].

As example we use here the same high-level functional categories as in previous work [23].

bN *<u>b</u>inding of <u>n</u>ucleic acids*: GO:0003676 at superfamily level.
bP *<u>b</u>inding of <u>p</u>roteins* with potential nuclear localization: GO:0005515 superfamily level.
rC *<u>r</u>egulation of <u>c</u>hromatin* GO:0016568 at superfamily level.
rB *<u>r</u>egulation of <u>b</u>inding*: GO:0051098 at superfamily level.
rE *<u>r</u>egulators of <u>e</u>nzymatic activity:* GO:0050790 at superfamily level.
mS *<u>m</u>etabolism of <u>s</u>accharides:* GO:0005976 at superfamily level.

The four functional groups bN, bP, rC, and rB encapsulate major modes of regulation. Both bN and bP play an important role for gene regulation by transcription factors and are among the most abundant GO classes, while rC focuses on chromatin-based epigenetic regulation. We have shown in [23] that rC group correlates well with the hand-picked collection of domain models that can act as readers, writers, and erasers of histone modification [25]. The domain groups rE and mS were intended as a form of controls that *a priori* we did not expect to correlate in a particular way with either nucleic acid or protein binding domains (bN, bP).

From the co-occurrences of domains in predicted proteins and the map of domains to functional (GO-)classes it is straightforward to obtain the number *n*(*C*, *D*) of co-occurrences of the functional classes. As in [23] we correct *n*(*C*, *D*) for the fact that the same domain *x* can be a member of both *C* and *D* by counting these cases with a weight of 1/2.

### 2.4 Co-occurrence Analysis

For each of the 18 species, we separately evaluated the number of domain co-occurrences and the number of genes in which two domain types *x* and *y* co-occur. Here *x* and *y* can be either individual domains, sets of domains belonging to the same superfamily, or the collections of domains compiled into functional classes according to their GO annotations. Denote by *n_x_* the total number of annotated domains belonging to group *x*. The simplest estimate for the expected number of domain co-occurrences is *E*(*x*, *y*) = *n_x_n_y_*/*n_g_*, where *n_g_* is the number genes in the genome under consideration. As discussed in [23] this estimate does not account for biases arising from the non-uniform distribution of domains over genes. Let *n_d_*(*i*) be the number of domains predicted for protein *i*, and let *n_d_* = Σ_*i*_ *n_d_*(*i*) be the total number of domains. Then the number of *x*-domains that occur in genes that also contain a *y*-domain can be estimated as

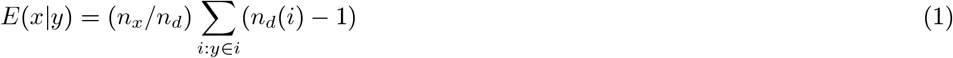

where the sum runs over all genes *i* that contain a domain belonging to group *y*. We obtain an alternative estimate by exchanging *x* and *y* in equ.(1).

We compared these expectations with the number of empirically observed co-occurrences *n*(*x*, *y*). We speak of *co-occurrence* of domain families or groups *x* and *y* if *n*(*x*, *y*) ≫ max{*E*(*x*|*y*), *E*(*y*|*x*)} and of *avoidance* if *n*(*x*, *y*) ≪ min{*E*(*x*|*y*), *E*(*y*|*x*)} The statistical significance of an observed difference between *n*(*x*, *y*) and the values of max{*E*(*x*|*y*), *E*(*y*|*x*)} and max{*E*(*x*|*y*), *E*(*y*|*x*)}, respectively, is determined under the assumption that *n*(*x*, *y*) is drawn from a Poisson distribution.

## 3 Results and Discussion

The comparison of the `AUGUSTUS` gene prediction results and `RefSeq` gene inventories agrees rather well in some species, while in others there are substantial differences, depending on the various degree of completeness of the gene annotation, Fig. 2. Since we are interested primarily in the distributions of protein domains we also compared `RefSeq` data with gene predictions restricted to only those genes in which at least one `Pfam` domain was annotated. For most species this improves the congruence between the gene sets. In a few cases, however, the differences persist, as in the case of *Trypanosoma* and Human, Fig. 2. In *Trypanosoma*, most of the difference is explained by annotated `RefSeq` proteins without recognizable domains. In human, the discrepancy is in part explained by RefSeq isoforms and in part by `AUGUSTUS` prediction without domains.

Among the predictions with annotated domains, we find e.g. for *Leishmania, Tetrahymena*, and *Plasmodium* that both the online and the offline trained gene predictions have a much larger coverage than the RefSeq data. For *Trichomonas* and *Giardia*, the situation is reversed. This can probably be explained in part by the large number of paralogs and possible pseudogenes included in RefSeq in *Trichomonas*, but also indicated as lack of sensitivity of the gene predictor for the two parabasalids with their extremely intron-poor genomes. At the domain level, `AUGUSTUS` and RefSeq agree nearly perfectly e.g. in human in *Naegleria*. In general, the RefSeq entries missed by the gene predictor are frequently putative pseudogenes and ORFs lacking further annotation. Since the `AUGUSTUS` ‘online’ predictions overall yield the most inclusive data set, these predictions are used below for all statistical analysis of domains compositions.

In general, we observe very little variation in the number of domains per protein. A significant increase is found in human and fruitfly only. It is unclear, however, whether this a true effect or an artifact arising from a bias in `Pfam` database. In [11], a difference in the complexity of chromatin proteins between Diplomonads and Dicristates on the one hand, and Alveolates and Stramenopiles on the other hand. Our data do not show such a systematic difference for proteins containing an rC domain.

In Figure 3 we observe a systematic avoidance of functionally distinct GO-classes of protein domains. Satisfactorily, the patterns obtained from `Pfam` and `SUPERFAMILY` annotations are largely consistent. Not surprisingly, we find fewer significant relations in the `SUPERFAMILY` data due the much smaller number of domains.

**Figure 3.**
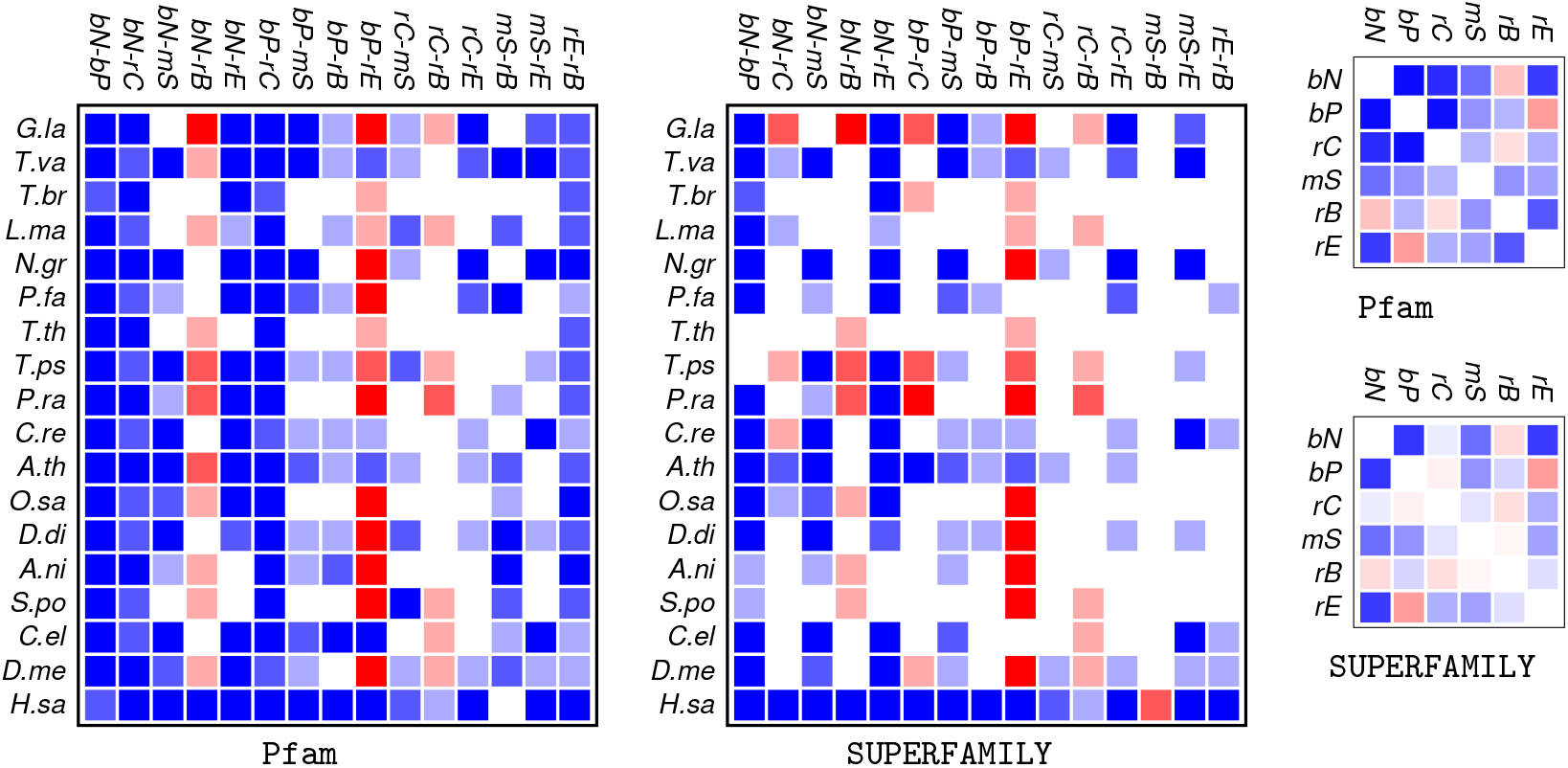
Summary of co-occurrences patterns of major functional classes of protein domains across the Eukaryotes. The estimated obtained from `Pfam`-domains (l.h.s.) are qualitatively consistent with those from `SUPERFAMILY`-domains (r.h.s). The top row shows the data separately for each species, the smaller panels below summarize the co-occurrence patterns across the 18 species. Blue rectangles indicate statistically significant avoidance between functional classes of protein domains, red indicates co-occurrence. The saturation of the color denotes the significance levels *p* < 0.001 (saturated color), 0.001 ≤ *p* < 0.01 (intermediate), and 0.01 ≤ *p* < 0.1 (pale). Entries that show neither avoidance or co-occurrence at a significance level of at least 10% remain white.

The main exceptions are the co-occurrences bN-rB, rC-rB, and bP-rE. The latter is not unexpected, since regulators of enzymatic activity (rE) can be expected to act by proteinprotein binding (bP). The positive correlations between nucleic acid binding domains (bN) and chromatin associated domains (rC) with domains involved in the regulation of binding deserved further investigation. It is consistent with intimate link of both DNA and RNA binding with chromatin regulation reported in [25].

The improved coverage and accuracy of the gene prediction procedure has a major impact on the observed domain co-occurrences. In an earlier study using the non-trainable genscan gene predictor we observed similarly wide-spread functional avoidance only for the large genomes of multicellular organisms [23]. At least a moderate positive correlation was found for most other genomes. In the light of the present data, i.e., a much larger set of annotated domains as well as a substantially improved set of underlying gene predictions, these co-occurrences are largely identified as artifacts.

## 4 Conclusion

The distribution of protein domains is an informative fingerprint of metabolic and regulatory capabilities of an organism. We have shown here that quantitative comparative analysis are possible based on predictions of trainable gene predictor such as `AUGUSTUS`. The training phase is necessary to overcome in particular artifacts introduced by peculiarities of the genome structure. Untrained tools such as `genscan`, for instance, have problems to recognize the protein boundaries in polycistronic transcripts of kinetoplastids, experience difficulties with extreme A/T contents, or lack sensitivity e.g. in very intron-poor genomes. Such effects are largely alleviated by species-specific training.

The second source of major ascertainment biases in the analysis of large scale evolutionary patterns of functional domains are the protein domain databases themselves. Recent studies reported the innovation of a large number of domain innovation events within both the green plants [12] and the animals [19]. The number of identified clade-specific domains must be expected to depend on the depths in which the clade is studied. The domain inventory is thus probably more complete in animals, fungi, and plants animals compared to most protozoan lineages. Large numbers unannotated domains of course undermine the analysis presented here since the lead to a systematic under-estimation of an organisms metabolic or regulatory capability, in particular since [12] also reported that the novel domains in stress response and developmental innovations. A more systematic survey of so-far undescribed protein domains thus constitutes a natural next step towards a comprehensive understanding of functional evolution in the eukaryotes.

Accurate domain inventories are not only of interest in their own right but also constitute an important source of phylogenetic information [36], in particular in “deep phylogeny” applications. The presence/absence patterns of protein domains were recently used for instance to place the Strepsiptera as a sister group of beetles in insect phylogeny [20]. Improved pipelines to estimate the protein domain content directly from genomic data thus have the potential to greatly facilitate phylogenomic investigations.

## Acknowledgment

We thanks Katharina Hoff (Greifswald) for technical assistance with the `AUGUSTUS` online gene prediction.

## Supplemental Data

http://www.bioinf.uni-leipzig.de/supplements/12-007

## References

1 G. Apic, J. Gough, and S. A. Teichmann. Domain combinations in archaeal, eubacterial and eukaryotic proteomes. J Mol Biol, 310:311–325, 2001.

2 S. L. Baldauf. An overview of the phylogeny and diversity of eukaryotes. J. Syst. Evol., 46:263–273, 2008.

3 E. Bornberg Bauer, A. K. Huylmans, and T. Sikosek. How do new proteins arise? Curr. Opin. Struct. Biol., 20:390–396, 2010.

4 M. Buljan and A. Bateman. The evolution of protein domain families. Biochem Soc Trans, 37:751–755, 2009.

5 C. Burge and S. Karlin. Prediction of complete gene structures in human genomic DNA. J. Mol. Biol., 268:78–94, 1997.

6 C. B. Burge and S. Karlin. Finding the genes in genomic DNA. Curr. Opin. Struct. Biol., 8:346–354, 1998.

7 D. A. de Lima Morais, H. Fang, O. J. Rackham, D. Wilson, R. Pethica, C. Chothia, and J. Gough. SUPERFAMILY 1.75 including a domain-centric gene ontology method. Nucleic Acids Res, 39:D427–D434, 2011.

8 I. A. Drinnenberg, D. E. Weinberg, K. T. Xie, J. P. Mower, K. H. Wolfe, G. R. Fink, and D. P. Bartel. RNAi in budding yeast. Science, 326:544–550, 2009.

9 S. R. Eddy. Accelerated profile HMM searches. PLoS Comp. Biol., 7:e1002195, 2011.

10 K. Forslund and E. L. L. Sonnhammer. Predicting protein function from domain content. Bioinformatics, 24:1681–1687, 2008.

11 L. M. Iyer, V. Anantharaman, M. Y. Wolf, and L. Aravind. Comparative genomics of transcription factors and chromatin proteins in parasitic protists and other eukaryotes. Int J Parasitol, 38:1–31, 2008.

12 A. R. Kersting, E. Bornberg-Bauer, A. D. Moore, and S. Grath. Dynamics and adaptive benefits of protein domain emergence and arrangements during plant genome evolution. Genome Biol Evol, 4:316–329, 2012.

13 K. M. Kim and G. Caetano-Anollés. The proteomic complexity and rise of the primordial ancestor of diversified life. BMC Evol Biol, 11:140, 2011.

14 E. Koonin, L. Aravind, and A. Kondrashov. The impact of comparative genomics on our understanding of evolution. Cell, 101:573–576, 2000.

15 F. Lu, H. Jiang, J. Ding, J. Mu, J. G. Valenzuela, J. M. C. Ribeiro, and X.-z. Su. cDNA sequences reveal considerable gene prediction inaccuracy in the *Plasmodium falciparum* genome. BMC Genomics, 8:255, 2007.

16 G. Melzer, R Theissen. MADS and more: transcription factors that shape the plant. Methods Mol Biol, 754:3–18, 2011.

17 S. Michaeli. Trans-splicing in trypanosomes: machinery and its impact on the parasite transcriptome. Future Microbiol., 6:459–474, 2011.

18 A. D. Moore, Å. K. Björklund, D. Ekman, E. Bornberg-Bauer, and A. Elofsson. Arrangements in the modular evolution of proteins. Trends Biochem. Sci., 33:444–451, 2008.

19 A. D. Moore and E. Bornberg-Bauer. The dynamics and evolutionary potential of domain loss and emergence. Mol Biol Evol, 29:787–796, 2012.

20 O. Niehuis, G. H. Hartig, S. Garth, H. Pohl, J. Lehmann, H. Tafer, A. Donath, V. Krauss, C. Eisenhardt, J. Hertel, M. Petersen, C. Mayer, K. Meusemann, R. S. Peters, P. F. Stadler, R. G. Beutel, E. Bornberg-Bauer, D. D. McKenna, and B. Misof. Genomic and morphological evidence converge to resolve the enigma of Strepsiptera. Current Biol., 2012.

21 T. Nilsson, M. Mann, R. Aebersold, J. R. Yates 3rd, A. Bairoch, and J. J. Bergeron. Mass spectrometry in high-throughput proteomics: ready for the big time. Nat Methods, 7:681–685, 2010.

22 C. A. Orengo and J. M. Thornton. Protein families and their evolution – a structural perspective. Annu Rev Biochem, 74:867–900, 2005.

23 A. A. Parikesit, P. F. Stadler, and J. Prohaska, Sonja. Evolution and quantitative comparison of genome-wide protein domain distributions. Genes, 2:912–924, 2011.

24 A. A. Parikesit, P. F. Stadler, and S. J. Prohaska. Quantitative comparison of genomic-wide protein domain distributions. In D. Schomburg and A. Grote, editors, German Conference on Bioinformatics 2010, volume P-173 of Lecture Notes in Informatics, pages 93–102, Bonn, 2010. Gesellschaft für Informatik.

25 S. J. Prohaska, P. F. Stadler, and D. C. Krakauer. Innovation in gene regulation: The case of chromatin computation. J. Theor. Biol., 265:27–44, 2010.

26 M. Punta, P. C. Coggill, R. Y. Eberhardt, J. Mistry, J. Tate, C. Boursnell, N. Pang, K. Forslund, G. Ceric, J. Clements, A. Heger, L. Holm, E. L. Sonnhammer, S. R. Eddy, A. Bateman, and R. D. Finn. The Pfam protein families database. Nucleic Acids Res, 40:D290–D301, 2012.

27 J. Schug, S. Diskin, J. Mazzarelli, B. P. Brunk, and C. J. Stoeckert Jr. Predicting Gene Ontology functions from ProDom and CDD protein domains. Genome Res, 12:648–655, 2002.

28 E. Shelest. Transcription factors in fungi. FEMS Microbiol Lett, 286:145–151, 2008.

29 M. Stanke. Lab session on gene prediction with AUGUSTUS, 2011. http://bioinf.uni-greifswald.de/augustus/binaries/tutorial/training.html.

30 M. Stanke, M. Diekhans, R. Baertsch, and D. Haussler. Using native and syntenically mapped cDNA alignments to improve *de novo* gene finding. Bioinformatics, 24:637–644, 2008.

31 M. Stanke, O. Schöffmann, B. Morgenstern, and S. Waack. Gene prediction in eukaryotes with a generalized hidden Markov model that uses hints from external sources. BMC Bioinformatics, 7:62, 2006.

32 M. Stanke and S. Waack. Gene prediction with a hidden Markov model and a new intron submodel. Bioinformatics, 19:ii215–ii225, 2003.

33 The Gene Ontology Consortium. Gene ontology: tool for the unification of biology. Nat Genet, 25:25–29, 2000.

34 S. Thomas, A. Green, N. R. Sturm, D. A. Campbell, and P. J. Myler. Histone acetylations mark origins of polycistronic transcription in *Leishmania major*. BMC Genomics, 10:152, 2009.

35 S. Yang and P. E. Bourne. The evolutionary history of protein domains viewed by species phylogeny. PLoS ONE, 4:e8378, 2009.

36 S. Yang, R. F. Doolittle, and P. E. Bourne. Phylogeny determined by protein domain content. Proc. Natl. Acad. Sci. USA, 102:373–378, 2005.

37 C. M. Zmasek and A. Godzik. Strong functional patterns in the evolution of eukaryotic genomes revealed by the reconstruction of ancestral protein domain repertoires. Genome Biol., 12:R4, 2011.

